# Integrated Transcriptomic and CRISPR Dependency Analysis Prioritizes a CDK1–AURKB Mitotic Vulnerability Axis in Diffuse Intrinsic Pontine Glioma

**DOI:** 10.64898/2026.08.30.748149

**Authors:** Arnav Rajeev, Soumith Veeragoni, Anish Sundar

**Affiliations:** Austin Community College

## Abstract

Diffuse intrinsic pontine glioma (DIPG), now classified within diffuse midline glioma, H3K27-altered, remains a lethal pediatric brainstem tumor with limited therapeutic options. Here, we integrated public DIPG transcriptomic datasets, protein–protein interaction modeling, functional enrichment, immune deconvolution, survival analysis, and DepMap CRISPR dependency data to nominate candidate mitotic vulnerabilities. Differential expression analysis comparing 27 DIPG tumors with 6 brainstem low-grade glioma comparator samples identified a proliferative transcriptional program enriched for chromosome segregation, nuclear division, and cell-cycle pathways. Network analysis prioritized a compact mitotic hub module containing CDK1, AURKB, TOP2A, CDC20, CDCA8, and related G2/M regulators. CIBERSORT analysis of an independent DIPG cohort inferred low cytotoxic T-cell signal, consistent with an immune-cold phenotype, although immune-cell fractions require orthogonal validation. Survival analysis showed that neither inferred immune scores nor a composite mitotic hub score significantly stratified overall survival. DepMap CRISPR gene-effect data nominated CDK1, AURKB, TOP2A, and BIRC5 as candidate dependencies across brain tumor models. These findings provide a computational framework for prioritizing mitotic vulnerabilities in DIPG and support experimental validation in disease-relevant models.

## 1 Introduction

The central nervous system relies on glial cells to sustain a stable environment for neurons, providing structural support, metabolic regulation, and immune surveillance throughout the brain and spinal cord.[1, 2] Well-established glial roles include maintaining the ionic milieu of nerve cells, modulating the rate of nerve signal propagation, controlling the uptake of neurotransmitters, and aiding in recovery from neural injury.[1, 2] When glial cells undergo malignant transformation, the resulting tumors, collectively known as gliomas, account for the majority of primary brain malignancies, representing about 80% of all malignant brain tumors.[3] Gliomas are classified according to cell of origin, differentiation, and malignancy grade, with the WHO grading system dividing them into four grades ranging from least to most aggressive.[4]

Among glioma subtypes, astrocytomas, tumors arising from star-shaped glial cells called astrocytes, are the most prevalent. Astrocytomas are classified by the WHO from grade 1 to grade 4, with grades 1 and 2 considered low-grade and grades 3 and 4 considered high-grade.[5] Low-grade tumors such as pilocytic astrocytoma tend to grow slowly and, in some cases, are not considered cancerous; high-grade tumors spread rapidly within brain tissue and carry significantly worse prognoses.[5]

At the most severe end of this spectrum lies diffuse intrinsic pontine glioma (DIPG), a pediatric high-grade astrocytoma that originates in the pons of the brainstem.[6, 7] DIPG is an aggressive and universally fatal pediatric cancer originating from glial cells surrounding and safeguarding neurons in the brain, predominantly affecting children in the age groups 0–4 and 5–9 years.[6] The 2021 WHO CNS Classification unified all cases of pediatric diffuse midline gliomas in which an H3K27 alteration was found, formally designating the tumor type as “Diffuse Midline Glioma, H3K27-altered” within the pediatric-type diffuse high-grade glioma family.[4, 7] This reclassification represented a major conceptual shift, from understanding DIPG purely by where it grows to defining it by what it is at the molecular level. The H3K27M mutation causes a global reduction in levels of H3 lysine 27 trimethylation, ultimately leading to epigenetic dysregulation of cellular processes through inactivation of the Polycomb Repressive Complex 2.[8]

The location of DIPG within the pons makes it uniquely destructive. The pons governs critical involuntary functions including breathing, heart rate, and blood pressure, as well as sensory and motor processes such as vision, hearing, and balance. These tumors often affect deep midline brain structures in young children, and patients commonly present with serious symptoms such as headache, vomiting, nerve palsy, and ataxia.[9, 10] What further distinguishes DIPG from other high-grade tumors is its diffuse growth pattern: rather than forming a discrete, removable mass, tumor cells infiltrate and intermingle with healthy brainstem tissue, rendering surgical resection effectively impossible without fatal damage to surrounding structures.[7, 12]

The clinical consequences are profound. Without treatment, most children survive only around six months following diagnosis; with radiation therapy, median survival extends to roughly nine to eleven months, and fewer than 10% of patients survive beyond two years.[7, 10] The mortality rate is exceptionally high, with a mere 1% five-year survival rate for newly diagnosed patients.[7] No chemotherapy regimen has demonstrated a meaningful impact on overall survival, and recurrence is virtually universal.[10] Despite significant advances in understanding DIPG at the molecular level, including the identification of recurrent H3K27M, ACVR1, TP53, and PDGFRA alterations, prognosis has remained essentially unchanged, making DIPG one of the most pressing unsolved problems in pediatric oncology.[7, 11, 12]

Recent high-dimensional profiling studies have begun to reveal the complex transcriptional and epigenetic landscapes of H3K27M-altered diffuse midline gliomas, identifying proliferative OPC-like tumor cell populations as major drivers of disease and highlighting potential therapeutic vulnerabilities.[13, 14] At the same time, multiple groups have shown that pediatric brain tumors, including DIPG, often reside in uniquely immune-cold microenvironments characterized by sparse T cell infiltration and abundant immunosuppressive myeloid and stromal populations, raising questions about the potential efficacy of conventional immunotherapy.[40]

In this context, there is a critical need to systematically identify tumor-intrinsic dependencies in DIPG and diffuse midline glioma that might be exploited therapeutically, alongside or independent of immune modulation. Here, we integrate differential expression analysis, protein–protein interaction (PPI) network modeling, functional enrichment, immune deconvolution, survival analysis, and genome-scale CRISPR dependency data to: (i) define a compact mitotic hub gene module in DIPG; (ii) characterize the associated immune microenvironment; and (iii) prioritize candidate hub genes with strong essentiality profiles in brain tumor models.

## 2 Materials and Methods

### 2.1 Data Acquisition

Two publicly available gene expression datasets were retrieved from the NCBI Gene Expression Omnibus (GEO).[15] The primary dataset (GSE26576) was generated on the Affymetrix Human Genome U133 Plus 2.0 platform (GPL570) and comprised 27 DIPG tumor samples (24 postmortem, 3 pretreatment biopsies), 6 brainstem low-grade glioma (LGG) samples, and 2 normal pediatric brainstem samples.[16] Two additional GBM/HGG samples present in the series (GSM488003, GSM488008) were profiled on a different array platform (GPL6801) and were excluded from all analyses.[16] Because only two true normal brainstem samples were available, the primary differential-expression analysis compared DIPG tumors (*n* = 27) against brainstem LGG as a within-brainstem comparator group (*n* = 6).[39] This design identifies transcriptional features that distinguish high-grade from low-grade brainstem glioma rather than tumor versus normal tissue, consistent with approaches used in prior DIPG expression studies, and results should be interpreted in that context.[18, 39]

DIPG samples included both postmortem (*n* = 24) and pretreatment biopsy specimens (*n* = 3); given the limited sample size, pretreatment and postmortem samples were analyzed together, though treatment-related transcriptional changes in postmortem tissue cannot be fully excluded.[39, 41]

### 2.2 Data Preprocessing and Normalization

Processed GSE26576 series-matrix expression values were retrieved from GEO using the GEOquery package.[20] Expression values were inspected to determine whether log_2_ transformation was required. Probe-to-gene symbol mapping for the GPL570 platform was performed using hgu133plus2.db. Where multiple probes mapped to the same gene symbol, the probe with the highest mean expression across samples was retained. Genes in the bottom quartile of expression variance were removed before differential-expression analysis. Principal component analysis and sample-distance heatmaps were used to assess sample-level structure and obvious outliers; formal batch-effect correction was not performed for the primary GSE26576 comparison.

### 2.3 Differential Expression Analysis

Differential expression analysis between DIPG tumor samples and brainstem LGG comparator samples was performed using the limma package.[21] A linear model was fitted using lmFit(), followed by empirical Bayes moderation via eBayes(). Genes were considered differentially expressed at an adjusted p-value threshold of *<* 0.05 (Benjamini–Hochberg correction) and | log_2_ FC*| >* 1. Volcano plots were generated using ggplot2 and ggrepel; genes passing the adjusted p-value and fold-change thresholds were highlighted.[22, 23] Differentially expressed genes were therefore defined as those significantly upregulated or downregulated in DIPG relative to brainstem LGG comparator samples;[21, 39] genes overexpressed in DIPG versus LGG represent candidates for DIPG-specific transcriptional activation rather than pan-tumor versus normal differences.[18, 19]

### 2.4 Protein–Protein Interaction Network Construction

Differentially expressed genes were mapped to the STRING database (v11.5) using the STRINGdb R package with a minimum interaction score threshold of 400.[24] Of 2,134 significant gene-level DEGs, 1,870 were successfully mapped to STRING, yielding a network with 1,867 nodes and 18,521 edges after filtering to within-set interactions and simplifying duplicate edges. A PPI network was constructed using igraph and visualized with ggraph. Network communities were identified using the Louvain modularity optimization algorithm. Hub genes were ranked by hub score (computed via hub score() in igraph) and node degree. The top 20 hub genes from the corrected STRING network were selected for downstream analysis.

### 2.5 Functional Enrichment Analysis

Gene Ontology Biological Process (GO BP) and KEGG pathway over-representation analyses were performed on the top 20 hub genes using clusterProfiler (v4.8), with Entrez IDs mapped via org.Hs.eg.db.[25] Over-representation results were filtered at adjusted *p <* 0.05.

### 2.6 Immune Microenvironment Deconvolution

Immune cell infiltration was estimated from the GSE50021 expression matrix using CIBERSORT with the LM22 signature matrix, which deconvolves 22 immune cell subtypes.[26] Samples were analyzed in absolute mode. Cell type fractions were visualized as stacked barplots ordered by CD8 T cell fraction. The top 10 most variable immune cell types were identified by inter-sample variance and visualized as boxplots. Spearman correlations between hub gene expression and immune cell fractions were computed across the 35 DIPG samples and visualized as a correlation heatmap.[40]

### 2.7 Survival Analysis

Overall survival data were available for all 35 DIPG samples in GSE50021.[17] Kaplan–Meier survival curves were generated using the survival and survminer R packages.[27, 28] Samples were stratified by immune cell presence/absence (non-zero threshold) and by median splits for CD8 T cells, NK cells, mast cells, and naive B cells. A composite mitotic hub gene score was computed per sample as the mean z-score of the top 20 hub genes from the GSE50021 expression matrix. Samples were dichotomized at the median mitotic score. Log-rank p-values are reported for all comparisons.

### 2.8 Druggability Assessment and CRISPR Dependency

The top 20 hub genes were manually cross-referenced against DrugBank, ChEMBL, and published literature for: (i) existence of small-molecule inhibitors, (ii) blood–brain barrier (BBB) penetrance data, and (iii) preclinical or clinical evidence in glioma or DIPG specifically.[7, 11, 12, 31, 32]

Genome-scale CRISPR gene-effect scores were retrieved from DepMap Public 24Q4 using CRISPRGeneEffect.csv and Model.csv. These gene-effect scores represent post-Chronos estimates of knockout fitness effects. Brain/CNS/glioma models were identified from Model.csv using Oncotree lineage, primary disease, subtype, and related metadata fields containing CNS, brain, glioma, glioblastoma, astrocytoma, oligodendroglioma, medulloblastoma, or meningioma terms.[33–35]

For each hub gene, gene-effect distributions were summarized in brain/CNS/glioma versus non-brain contexts, and pairwise Spearman correlations were computed among selected mitotic regulators (CDK1, AURKB, TOP2A, BIRC5, BUB1B, TTK, CCNB1, BUB1, DLGAP5).

## 3 Results

### 3.1 Sample Quality Control and Expression Landscape

PCA of normalized microarray expression values showed separation between DIPG tumor samples and brainstem LGG comparator samples along PC1, consistent with condition-associated transcriptomic structure (Figure 1).[16, 21, 39] Sample distance heatmap analysis showed broad sample-level structure and no obvious outlier samples, although formal batch-effect correction was not performed (Figure 2).[21]

**Figure 1:**
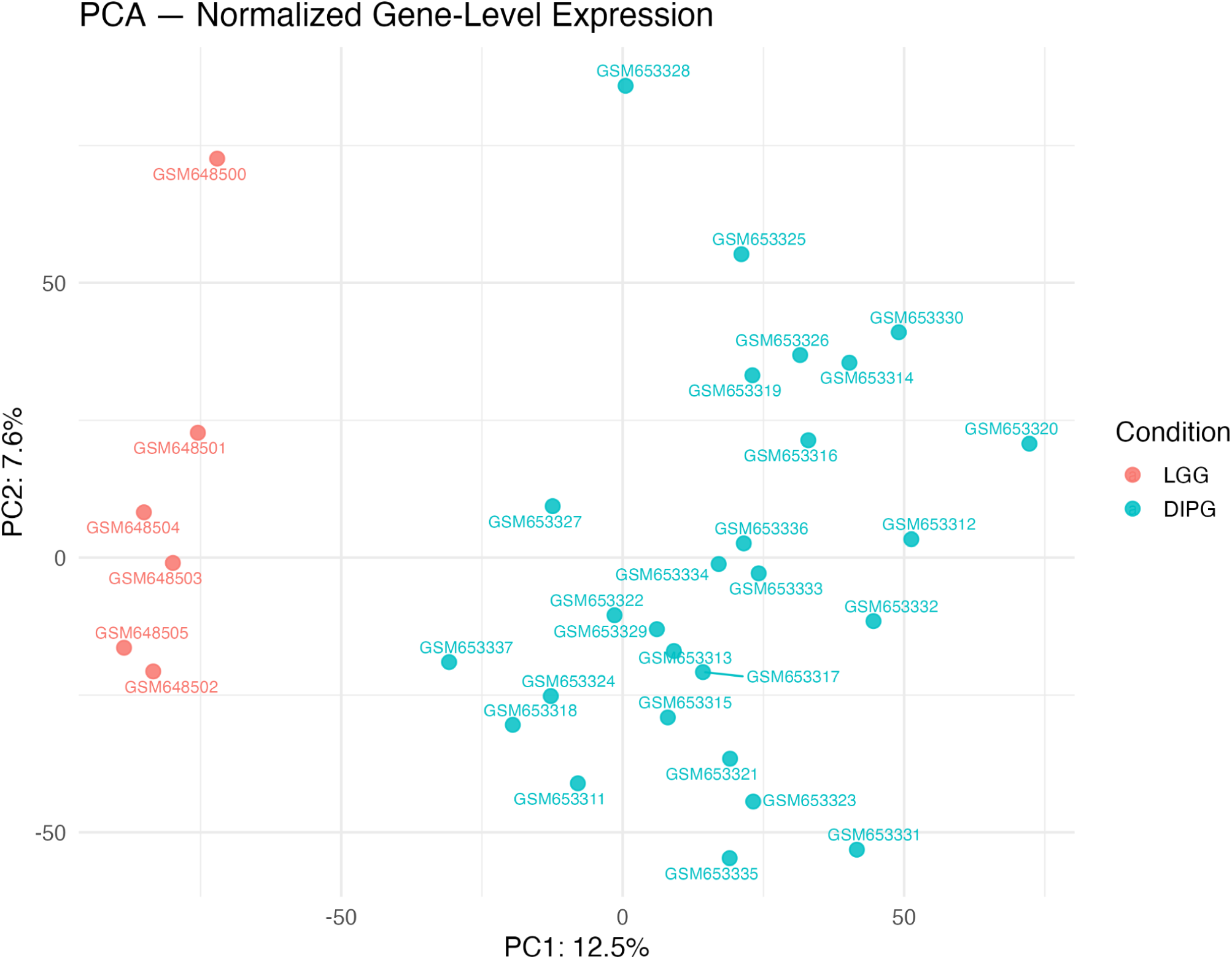
Principal Component Analysis of Normalized Microarray Expression Values. PCA of normalized microarray expression values across DIPG tumor samples and brainstem LGG comparator controls.[16, 21] Each point represents one sample, colored by condition. The plot shows separation between DIPG and brainstem LGG samples, supporting condition-associated transcriptomic structure while not by itself proving the absence of technical variation.

**Figure 2:**
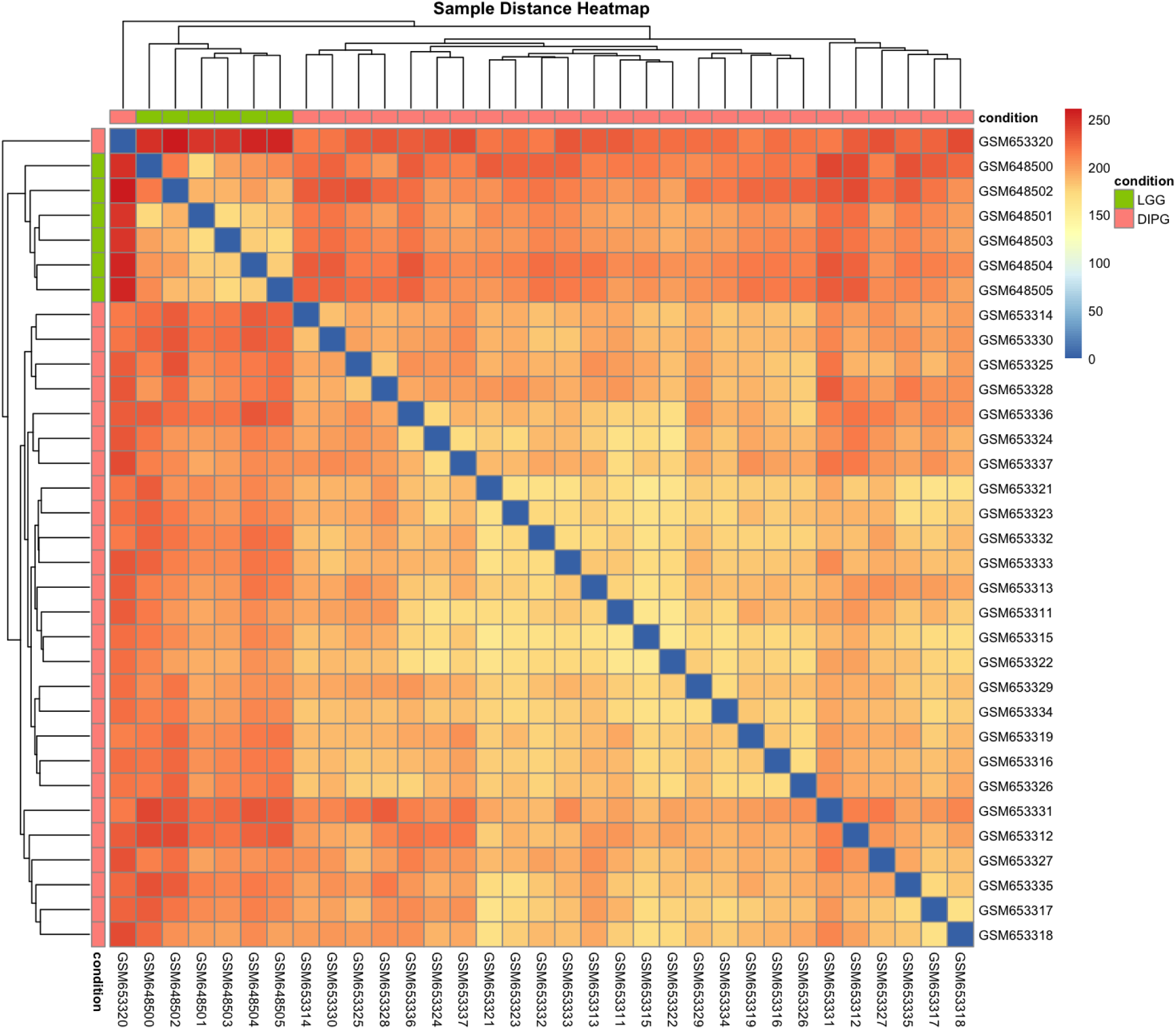
Sample Distance Heatmap. Euclidean distance matrix computed from normalized expression profiles across all samples.[16, 21] Color intensity represents pairwise similarity (dark = similar). The heatmap shows broad sample-level structure with no obvious outlier samples; formal batch-effect correction was not performed, and this figure supports general data quality but does not constitute proof of the absence of technical variation.

### 3.2 Differential Expression Analysis Identifies a Proliferative Transcriptomic Signature

Differential expression analysis comparing 27 DIPG tumors against 6 brainstem LGG samples identified 2,134 significant gene-level differentially expressed genes after probe-to-gene collapse and low-variance filtering (adjusted *p <* 0.05, | log_2_ FC*| >* 1), representing genes transcriptionally elevated or suppressed in high-grade relative to low-grade brainstem glioma.[18, 19, 21] Upregulated genes were predominantly associated with cell cycle progression and mitotic regulation, while downregulated genes were enriched for immune and inflammatory processes.[18, 19] The volcano plot (Figure 3) illustrates the distribution of differentially expressed genes, with top hub genes labeled.

**Figure 3:**
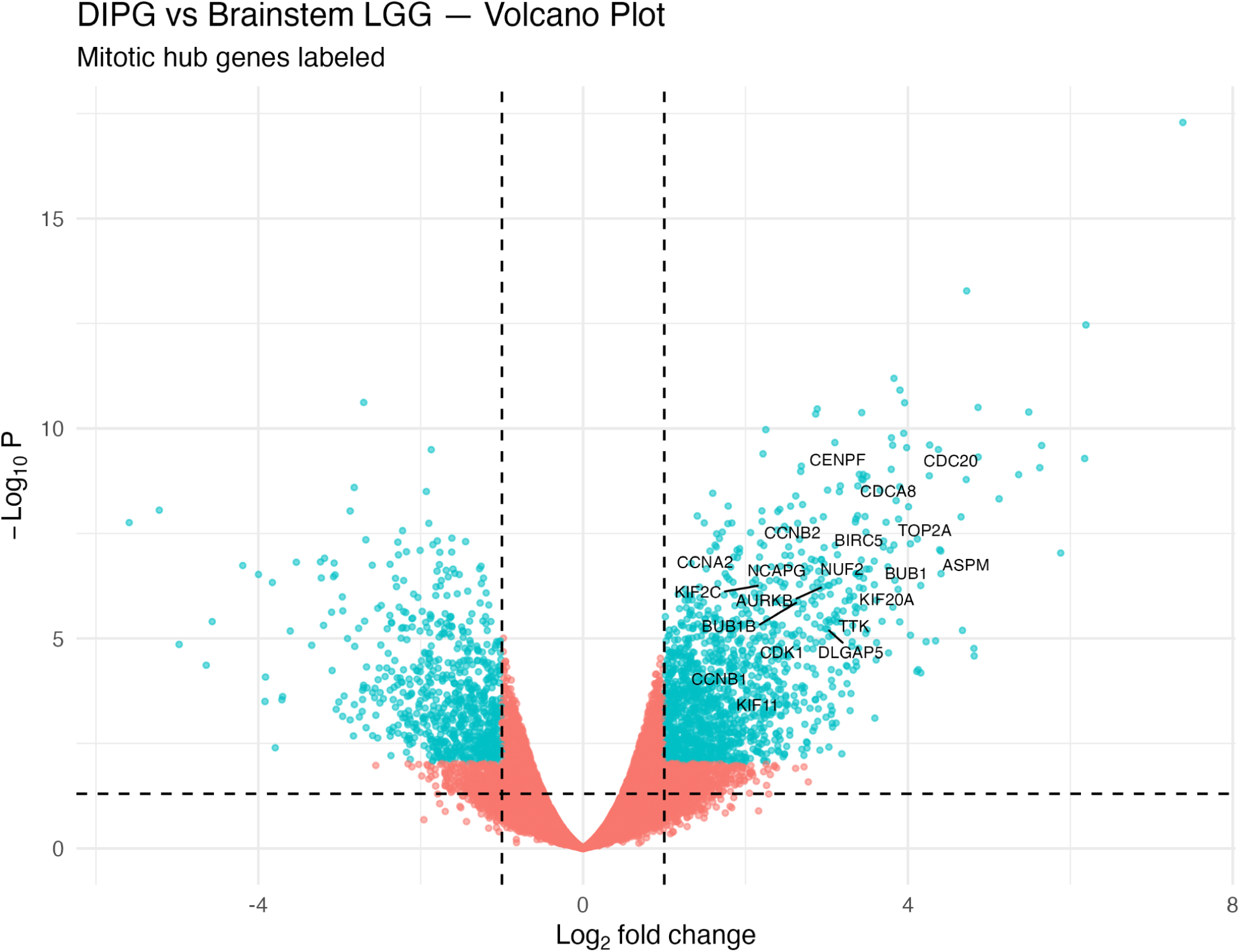
Volcano Plot of Differentially Expressed Genes in DIPG versus Brainstem LGG. Each point represents one gene, plotted by log_2_ fold change (x-axis) and *−* log_10_ nominal p-value (y-axis). Genes passing adjusted *p <* 0.05 and | log_2_ FC*| >* 1 are highlighted relative to brainstem LGG comparator samples (*n* = 6).[16, 39] Mitotic hub genes are labeled.

### 3.3 PPI Network Analysis Prioritizes a Mitotic Hub Gene Module

Of 2,134 significant gene-level DEGs, 1,870 mapped to STRING, yielding a corrected PPI network with 1,867 nodes and 18,521 edges.[24] Using hub score and degree, the corrected network prioritized a compact mitotic module containing CDK1, AURKB, TOP2A, CDC20, CDCA8, CCNA2, CCNB1, CCNB2, BUB1, BUB1B, TTK, KIF11, KIF20A, DLGAP5, ASPM, NCAPG, AURKA, CDC45, TPX2, and RRM2. Network community detection identified functional clusters enriched for spindle assembly checkpoint, cytokinesis, and chromosomal passenger complex components (Figure 4, 5). Expression heatmap analysis confirmed consistent and strong upregulation of all 20 hub genes across DIPG samples relative to brainstem LGG comparator samples (Figure 9).

**Figure 4:**
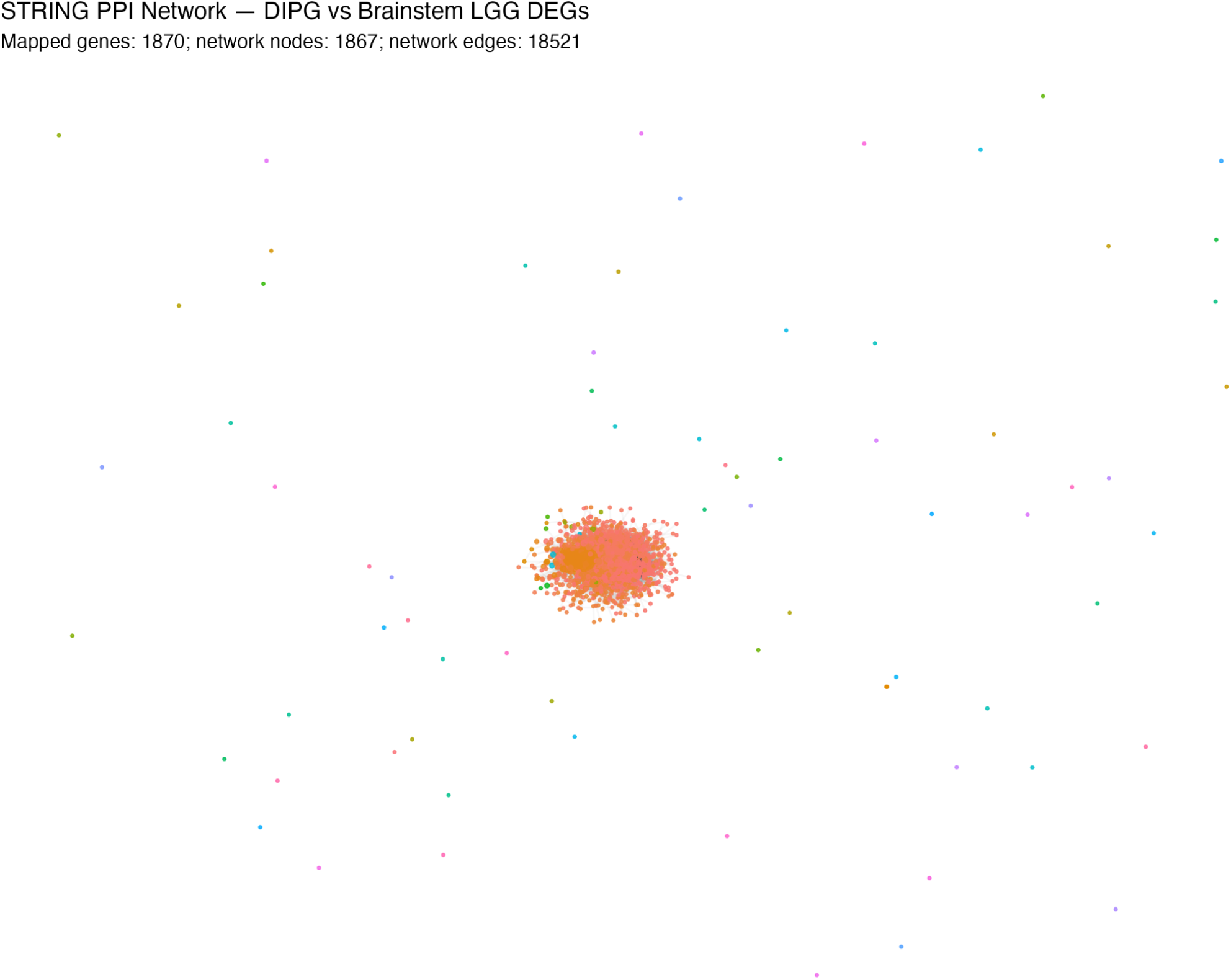
Protein–Protein Interaction Network of DIPG Differentially Expressed Genes. STRING-derived PPI network of 1,870 mapped gene-level DEGs, visualized using a Fruchterman–Reingold layout. The corrected network contained 1,867 nodes and 18,521 edges. Node size is proportional to hub score, node color represents Louvain community assignment, and edges represent protein interactions with a minimum confidence score of 400.

**Figure 5:**
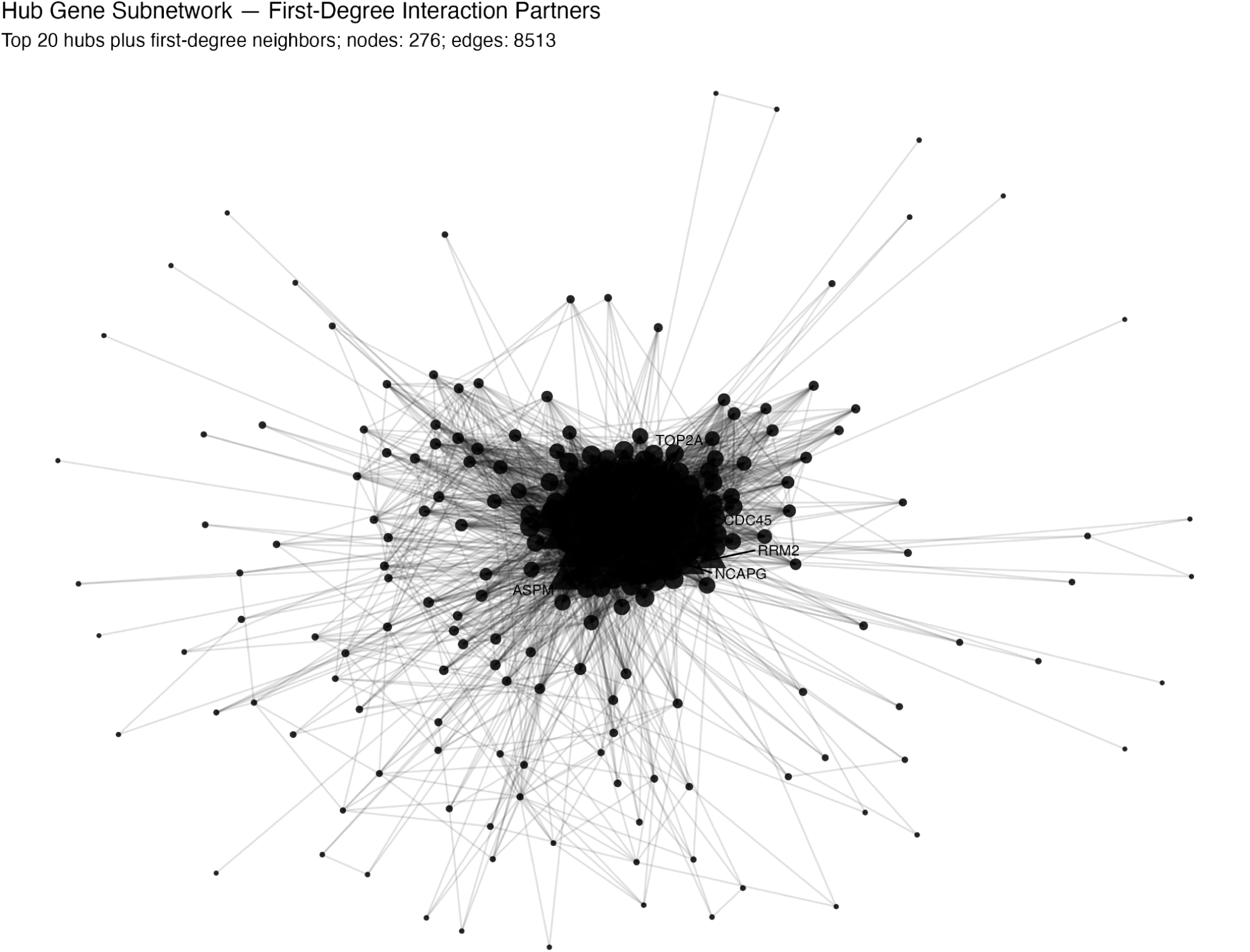
Hub Gene Subnetwork Showing First-Degree Interaction Partners. Subnetwork containing the corrected top 20 hub genes and their immediate first-degree interaction partners. The corrected subnetwork contained 276 nodes and 8,513 edges. Node size reflects hub score, and labeled nodes indicate selected high-ranking hubs within the mitotic module.

**Figure 6:**
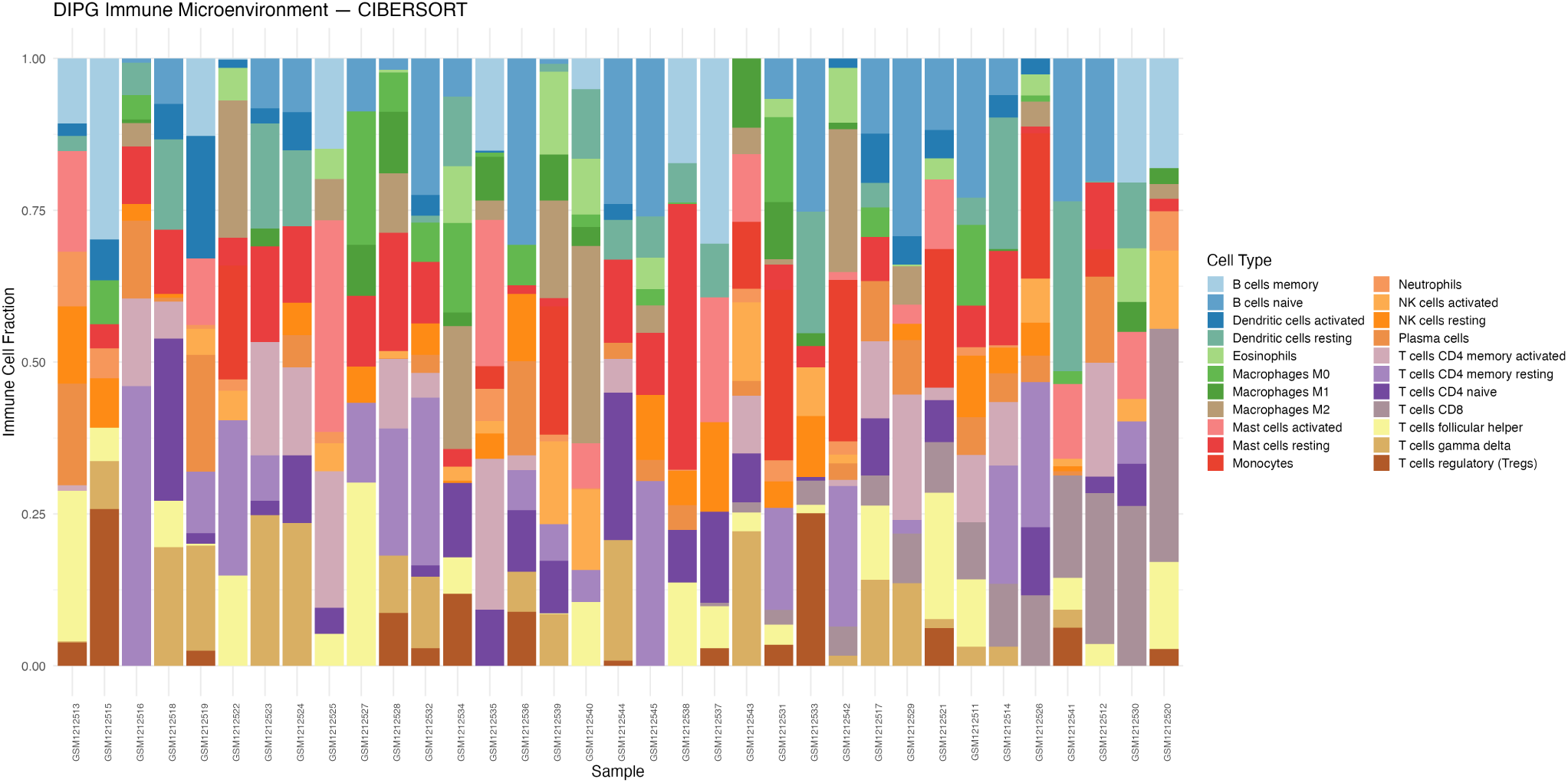
CIBERSORT Immune Cell Fraction Estimates Across DIPG Tumor Samples. Stacked barplot showing CIBERSORT-estimated proportions of 22 immune cell types across all 35 DIPG samples, ordered by CD8 T-cell fraction. Low inferred cytotoxic T-cell signal is consistent with an immune-cold phenotype, while specific LM22-derived mast-cell, B-cell, and myeloid fractions should be interpreted cautiously.

**Figure 7:**
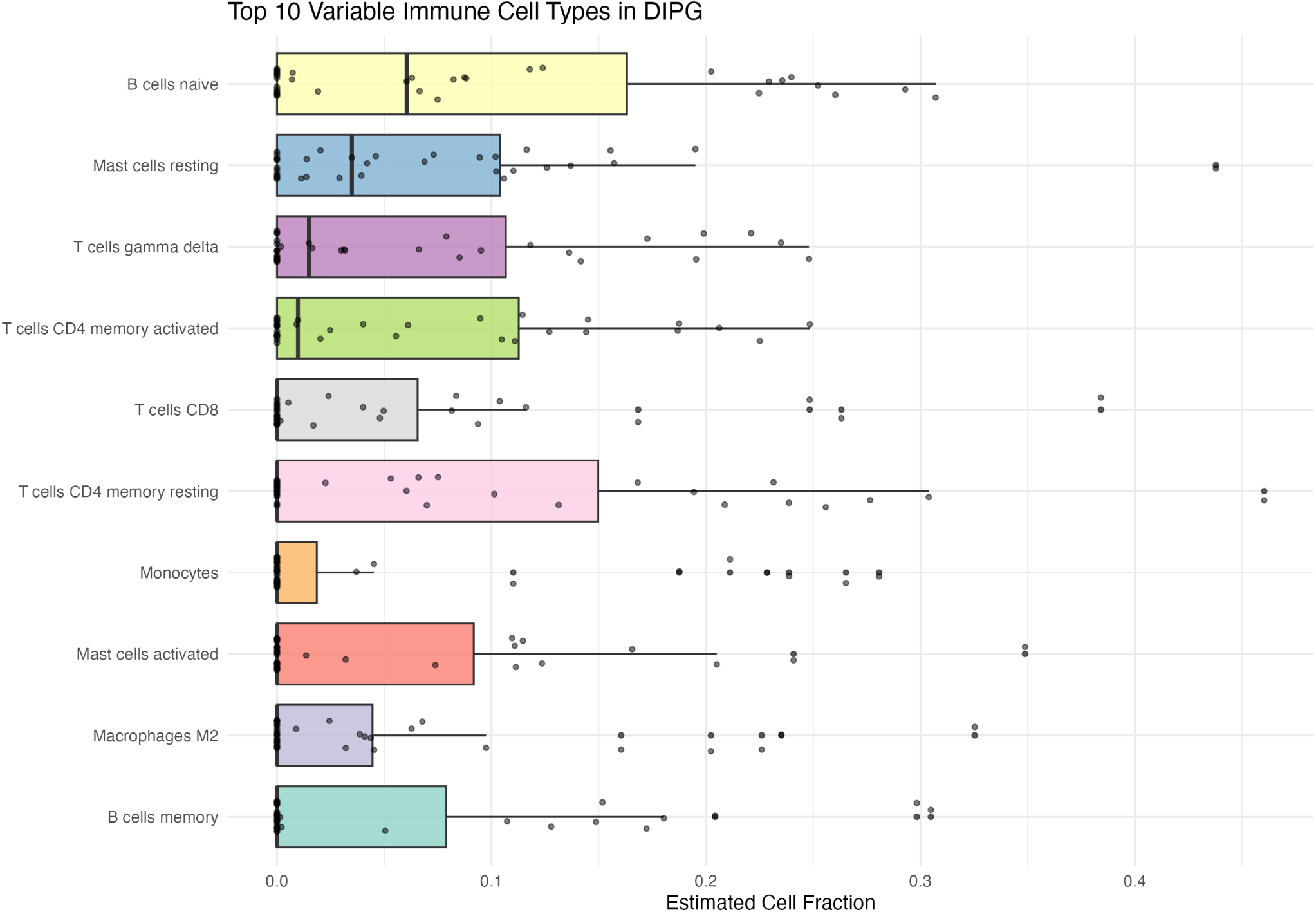
Distribution of the Top 10 Most Variable Immune Cell Types in DIPG. Boxplots with individual data points showing the inter-sample distribution of the 10 most variable inferred immune cell fractions, ordered by median fraction. Variation across LM22-estimated fractions suggests heterogeneity in immune-related expression signals across individual DIPG tumors.

**Figure 8:**
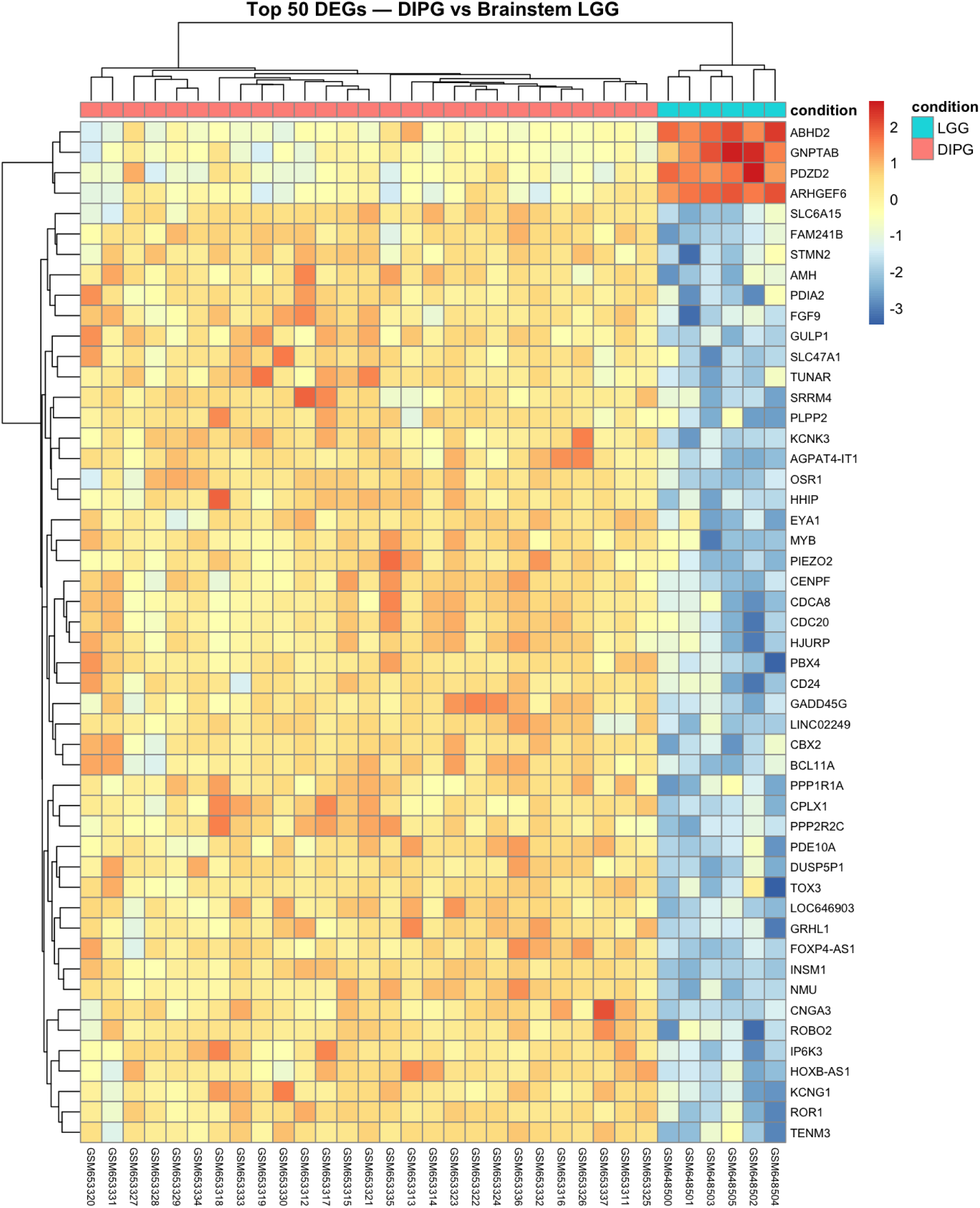
Heatmap of Top 50 Differentially Expressed Genes in DIPG versus Brainstem LGG. Z-score normalized expression of the 50 most significantly dysregulated genes across all samples, ranked by adjusted p-value.[18, 19] Columns are annotated by condition. Hierarchical clustering shows a clear expression shift between DIPG and brainstem LGG comparator samples, including upregulation of multiple mitotic regulators in the DIPG group.[18, 19]

**Figure 9:**
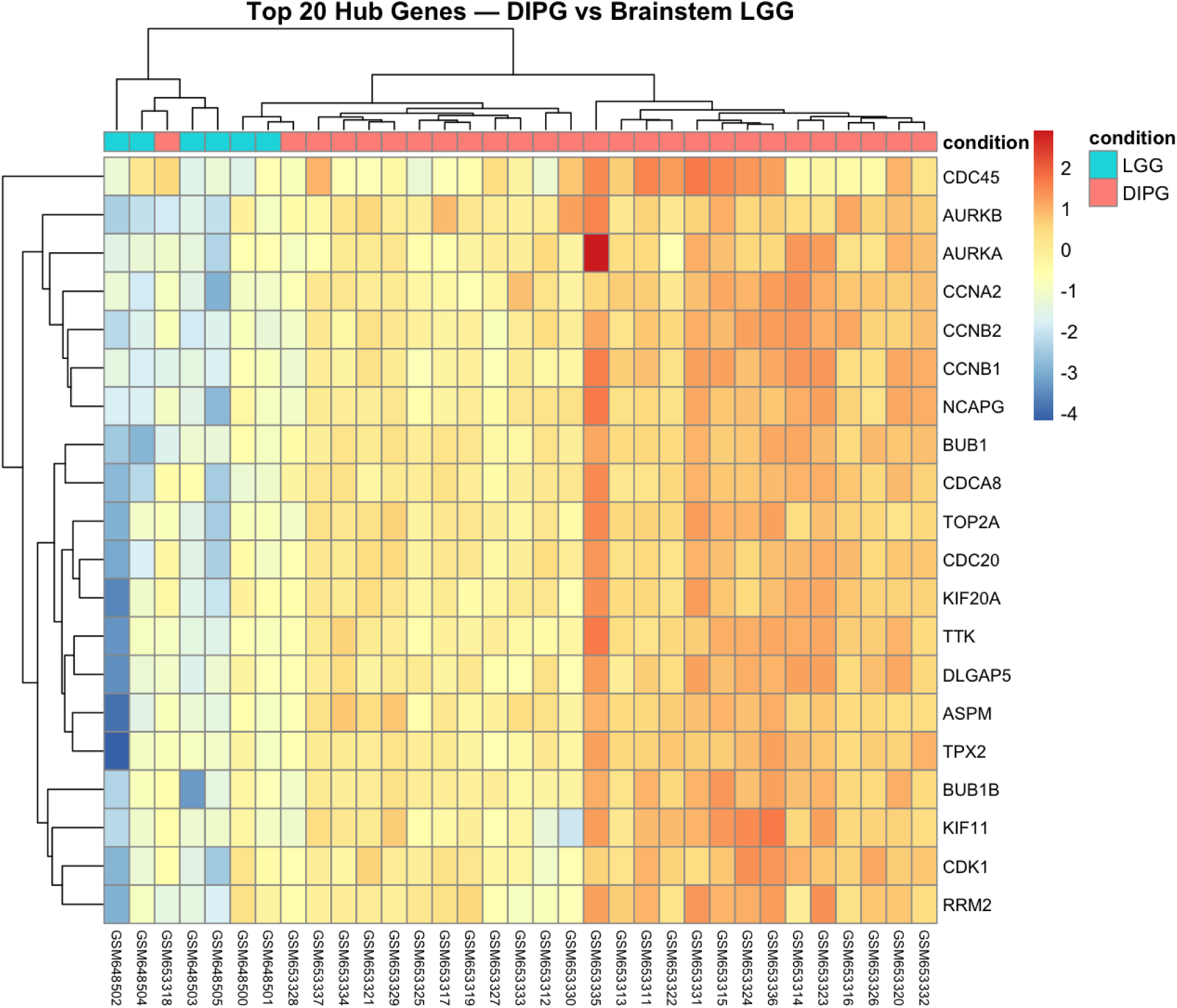
Expression Heatmap of Top 20 Hub Genes Across DIPG Samples. Z-score normalized expression of the 20 highest-ranked hub genes from the corrected STRING network across DIPG and brainstem LGG samples. Column annotations show condition. The corrected hub panel shows broad upregulation in DIPG relative to brainstem LGG comparator samples.

### 3.4 Functional Enrichment Confirms a Hyperactivated Mitotic Program

Over-representation analysis of the corrected hub genes yielded highly significant GO BP terms centered on nuclear chromosome segregation, chromosome segregation, nuclear division, sister chromatid segregation, and mitotic cell-cycle phase transition (Figure 10).[25] KEGG analysis confirmed cell cycle as the dominant pathway, with additional enrichment in progesterone-mediated oocyte maturation, oocyte meiosis, p53 signaling, and cellular senescence (Figure 11).[25]

**Figure 10:**
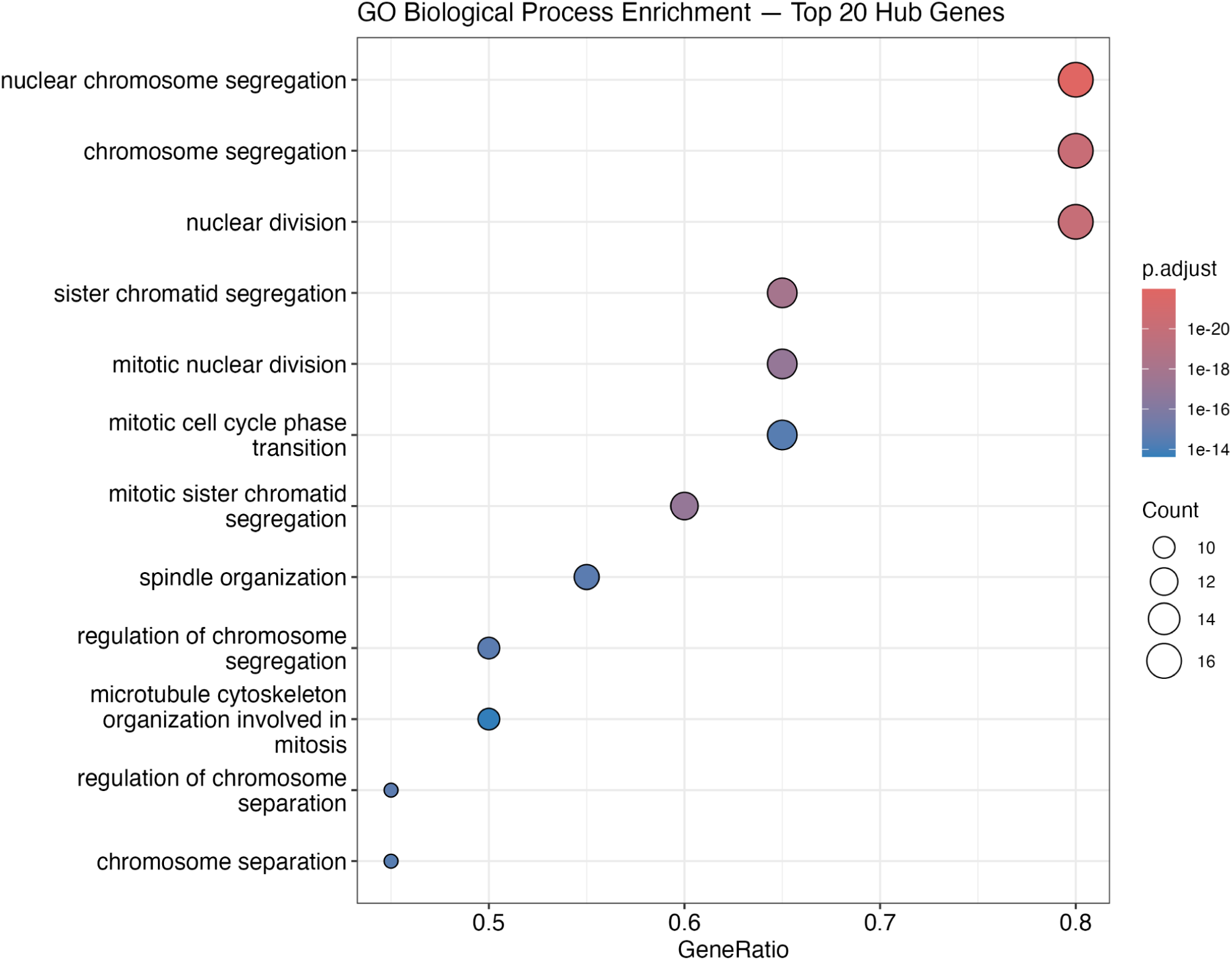
Gene Ontology Biological Process Over-Representation Analysis of Hub Genes. Dotplot showing the top enriched GO BP terms among the 20 hub genes. Dot size represents gene ratio; dot color represents adjusted p-value. All top terms relate to mitotic regulation and chromosome segregation.

**Figure 11:**
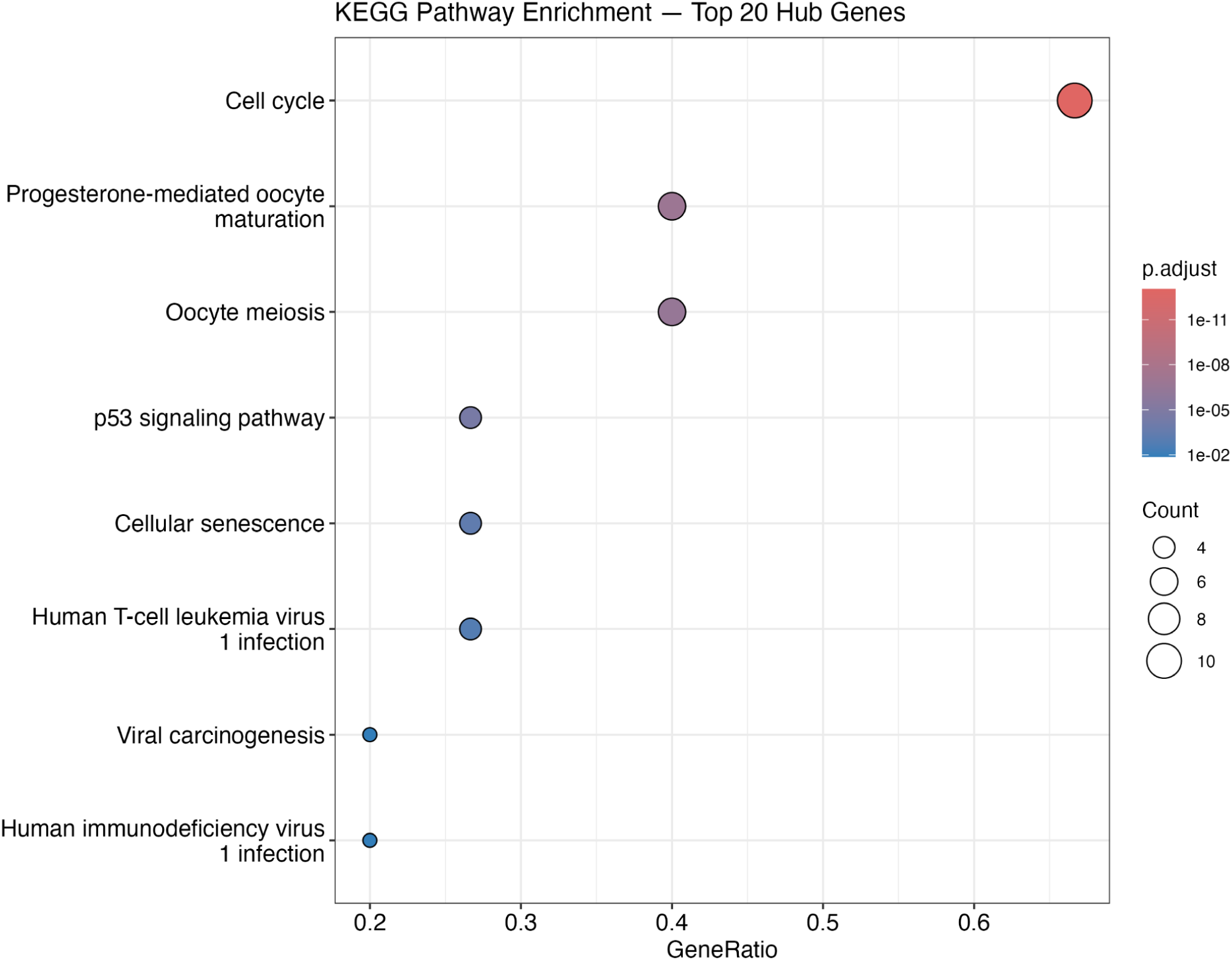
KEGG Pathway Over-Representation Analysis of Hub Genes. Dotplot of the top enriched KEGG pathways among hub genes. Cell cycle (hsa04110) is the most significantly enriched pathway, with additional enrichment in oocyte meiosis and p53 signaling.

### 3.5 Immune Deconvolution Reveals an Immune-Cold DIPG Microenvironment

CIBERSORT deconvolution of GSE50021 inferred low cytotoxic T-cell signal across most DIPG samples, consistent with an immune-cold phenotype.[17, 26] Because LM22-based deconvolution was not designed specifically for pediatric CNS tumors, inferred mast-cell, B-cell, and myeloid fractions should be interpreted cautiously and would require validation by orthogonal methods such as immunohistochemistry or single-cell sequencing.[26, 40] Correlation analysis between mitotic hub gene expression and immune fractions revealed predominantly negative associations, suggesting that hyperactive mitotic programs coexist with suppressed or altered local immune infiltration (Figure 12).

**Figure 12:**
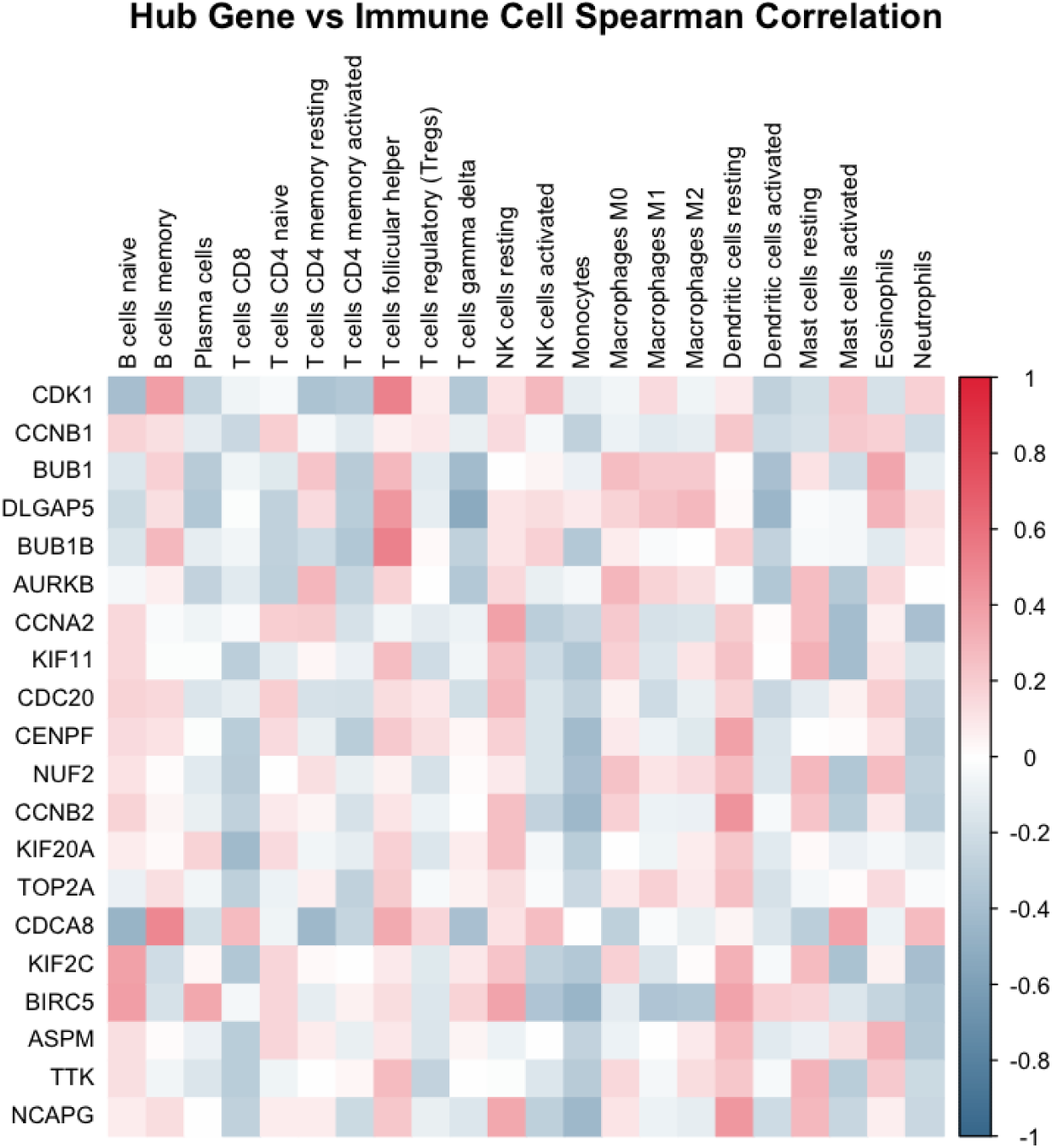
Correlation Between Hub Genes and Immune Cell Fractions. Spearman correlation heatmap between hub gene expression and selected CIBERSORT-inferred immune fractions in DIPG (GSE50021). Mitotic hub genes are generally anti-correlated with several inferred immune-cell signals, although these correlations require orthogonal validation.

### 3.6 Mitotic and Immune Program Scores Do Not Stratify DIPG Survival

We next assessed whether immune or mitotic programs carry prognostic information in DIPG. Using curated signatures for B cells, CD8 T cells, NK cells, and mast cells, we computed per-sample immune scores and stratified tumors by median score.[29, 30] For each lineage, high versus low immune score groups showed no significant difference in overall survival: B cell (log-rank *p* = 1.0), CD8 T cell (*p ≈* 0.67), NK cell (*p ≈* 0.79), and mast cell (non-significant), with no significant survival association observed in Kaplan–Meier analysis.[17]

We then evaluated the 20-gene mitotic hub module in GSE50021 by computing a composite mitotic score and splitting tumors at the median. Kaplan–Meier analysis similarly revealed no significant difference in OS between high and low mitotic groups (log-rank *p ≈* 0.87), with no significant survival association observed.[17] Univariate Cox models using standardized mitotic and immune scores yielded hazard ratios near 1 with wide confidence intervals, and a multivariable model including all five scores did not identify any independent prognostic effect. These findings indicate that, within a small and uniformly aggressive DIPG cohort, neither inferred immune infiltration nor mitotic activity alone is sufficient to stratify survival.

### 3.7 CDK1–AURKB Emerge as Candidate Mitotic Dependencies in Brain Tumor Models

Although mitotic and immune scores do not stratify survival, we hypothesized that the mitotic hub genes represent functional dependencies. DepMap CRISPR gene-effect data indicated that CDK1, AURKB, TOP2A, and BIRC5 are among the strongest candidate dependencies across 91 brain/CNS/glioma models, with mean gene-effect scores below *−*2 and nearly all models scoring below *−*1.0 for these genes (Figure 13).[33–35] BUB1B and TTK also showed strong essentiality, whereas CCNB1, BUB1, and DLGAP5 displayed somewhat weaker but frequent dependencies.

**Figure 13:**
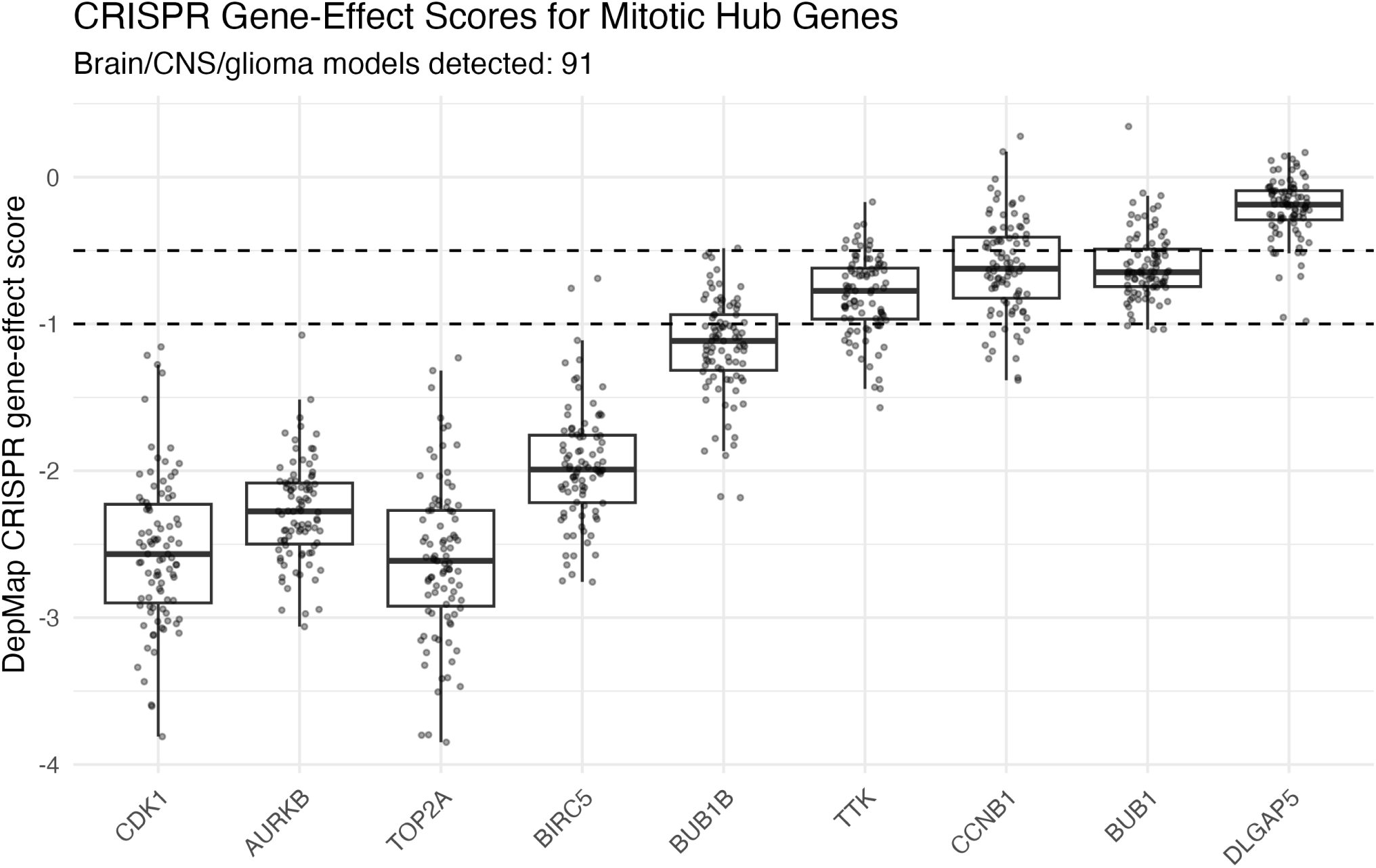
CRISPR Gene-Effect Scores for Mitotic Hub Genes in Brain Tumor Cell Lines. Boxplots showing gene-effect distributions for CDK1, AURKB, TOP2A, BIRC5, BUB1B, TTK, CCNB1, BUB1, and DLGAP5 across 91 brain/CNS/glioma models in DepMap. Dashed lines indicate dependency (*−*0.5) and strong essentiality (*−*1.0) thresholds. CDK1, AURKB, TOP2A, and BIRC5 show strong dependency profiles across most brain/CNS/glioma models.

Comparing gene-effect distributions in brain versus non-brain cell lines revealed that CDK1 and AURKB are broadly essential, with mean scores in brain tumors comparable to or more negative than those in other lineages (Figure 14).[34]

**Figure 14:**
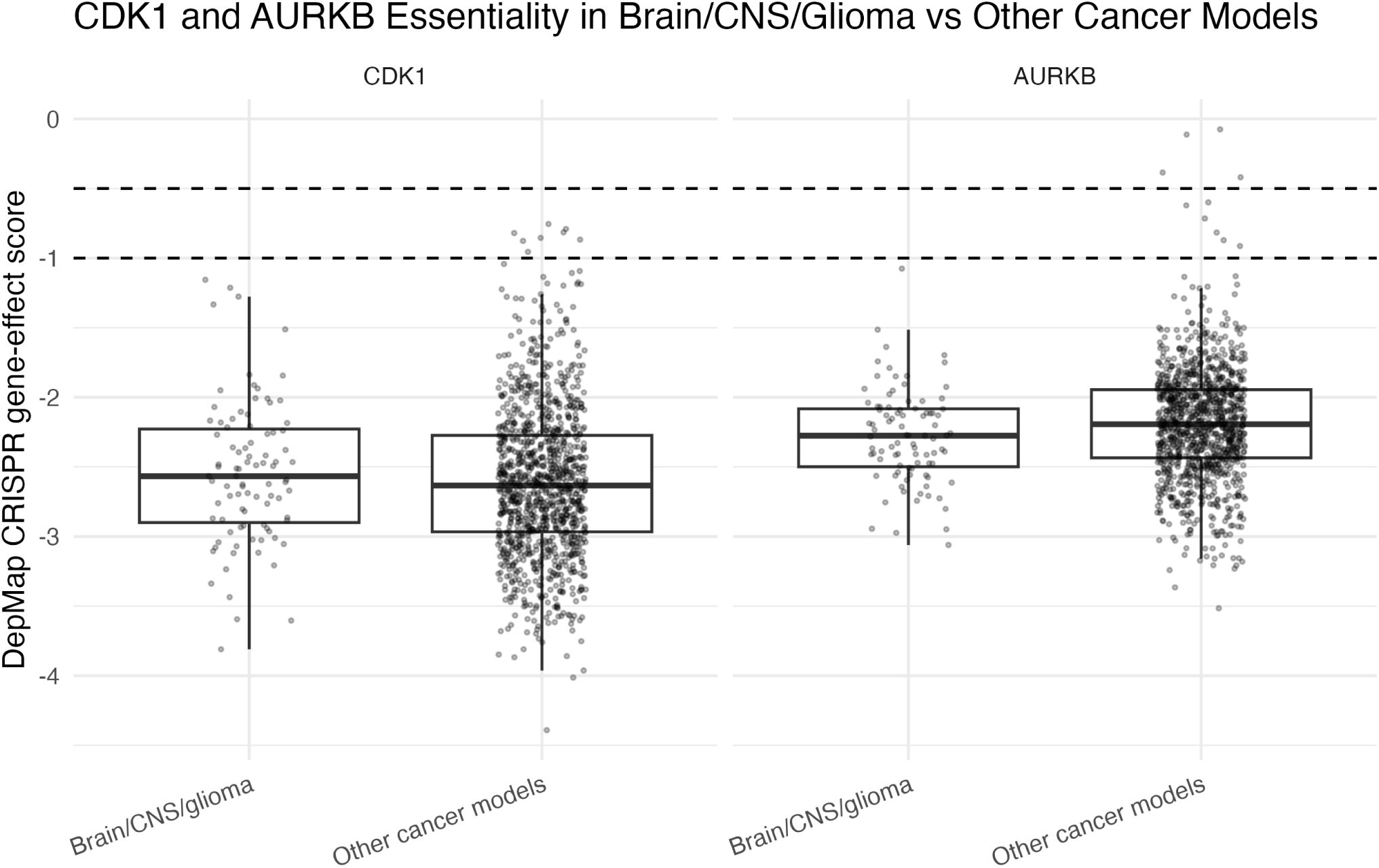
CDK1 and AURKB Essentiality in Brain versus Non-Brain Cancer Cell Lines. CRISPR gene-effect scores for CDK1 and AURKB in brain and non-brain cell lines from DepMap. Both genes show strong dependency profiles across contexts, underscoring their role as conserved mitotic dependencies that warrant disease-relevant validation in DIPG/DMG models.

Pairwise Spearman correlations among gene-effect scores for CDK1, AURKB, TOP2A, BIRC5, BUB1B, TTK, CCNB1, BUB1, and DLGAP5 revealed modest co-dependency structure among selected mitotic regulators, with correlation strength varying across genes (Figure 15).[33, 37] These results nominate CDK1–AURKB as candidate members of a broader G2/M dependency panel in brain tumor models, while emphasizing that functional validation remains necessary.

**Figure 15:**
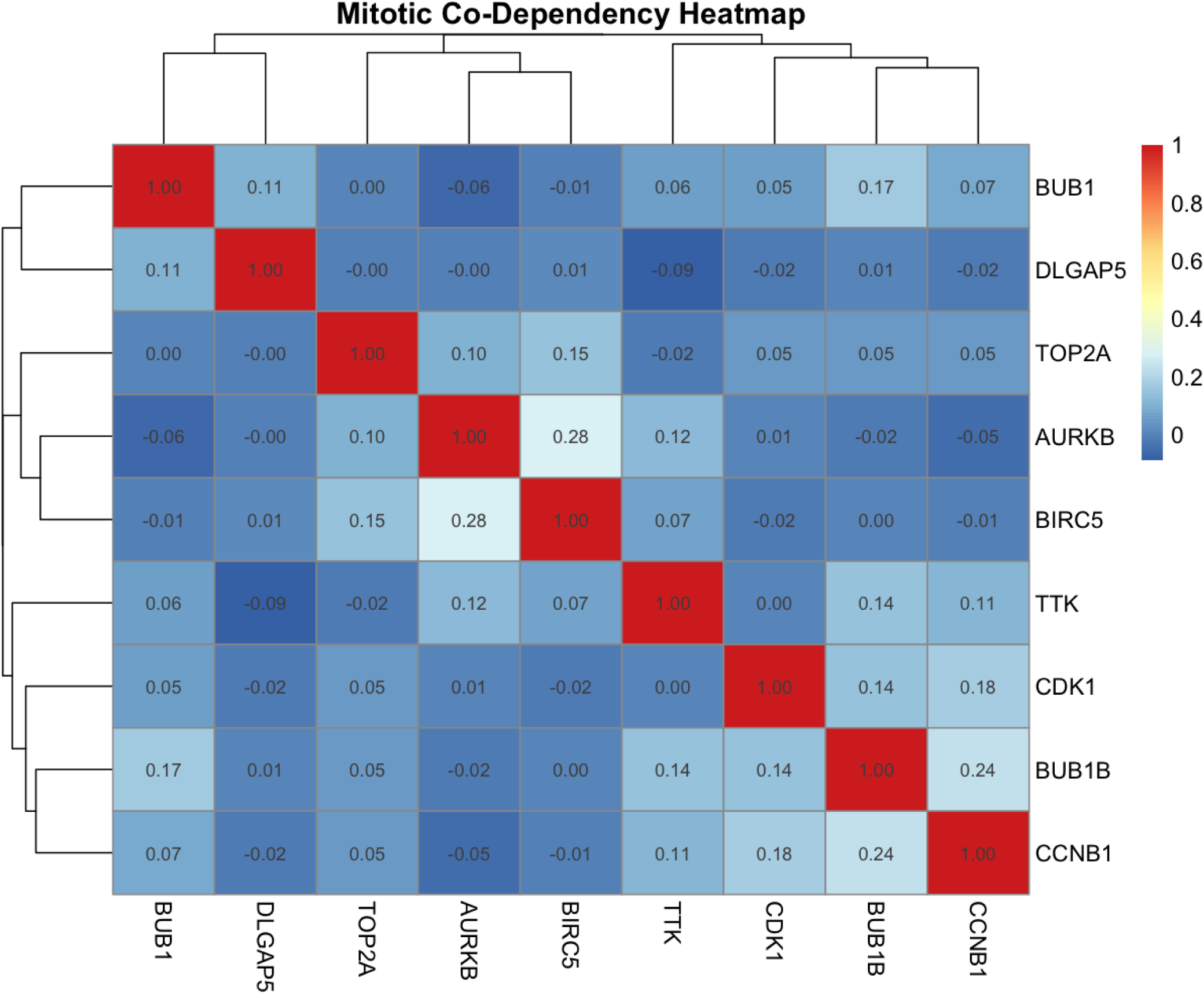
Mitotic Co-Dependency Heatmap. Spearman correlation heatmap of CRISPR gene-effect scores for mitotic hub genes across DepMap cell lines. The selected mitotic regulators show variable pairwise co-dependency correlations across DepMap cell lines.

## 4 Discussion

By integrating transcriptomic profiling, PPI network analysis, functional enrichment, immune deconvolution, survival modeling, and CRISPR dependency data, we support a model in which DIPG displays a strong mitotic transcriptional program within an immune-cold context.[7, 13, 14, 40] Our differential expression and network analyses highlight consistent overexpression of cell-cycle and mitotic regulators, with a corrected 20-gene hub module enriched for chromosome segregation and nuclear division. Immune deconvolution inferred low cytotoxic T-cell signal across most DIPG samples, aligning with prior descriptions of an immune-cold DIPG microenvironment, although specific LM22-derived cell-fraction estimates require validation by orthogonal approaches.[40] It is important to note that the primary DEG analysis compared DIPG against brainstem LGG rather than normal tissue; the identified mitotic hub genes therefore reflect transcriptional features that distinguish high-grade from low-grade brainstem glioma, and their absolute expression levels relative to normal brain remain to be established in larger matched datasets.[16, 18, 19, 39] Notably, neither immune scores nor a composite mitotic hub score significantly stratify overall survival in GSE50021, likely reflecting the uniformly poor prognosis of DIPG and limited power of small cohorts. Yet, DepMap data reveal that CDK1, AURKB, TOP2A, and BIRC5 are among the most strongly essential genes across brain tumor models, with broad dependency across models and variable co-dependency structure.[33–35] DIPG shows a strong mitotic transcriptional program, and DepMap CRISPR analysis nominates CDK1, AURKB, TOP2A, and BIRC5 as candidate mitotic dependencies in brain tumor models.[33–35] Because the DepMap panel contains few if any H3K27M-specific DIPG lines, experimental validation in patient-derived DIPG models—including neurospheres, organoids, and orthotopic xenografts—will be required to confirm these dependencies in a disease-relevant context.[7, 11, 41]

Future work should extend these analyses to multi-omic pediatric brain tumor atlases and single-cell datasets to localize the mitotic hub module to specific malignant cell states (e.g., OPC-like cycling compartments) and to preclinical DIPG models to experimentally validate CDK1–AURKB co-targeting strategies.[13, 14, 38] Nonetheless, the integrative pipeline presented here provides a rational framework for prioritizing mitotic vulnerabilities and complements ongoing efforts to modulate the immune microenvironment in diffuse midline glioma.

## Acknowledgements

The authors thank the patients and families who contributed samples to the GEO datasets used in this study (GSE26576, GSE50021).[16, 17] No external funding was received for this work.

## Data Availability

All expression data used in this study are publicly available through the NCBI Gene Expression Omnibus under accession numbers GSE26576 and GSE50021.[15–17] Genome-scale CRISPR dependency data were obtained from DepMap Public 24Q4. The corrected GSE26576 differential-expression analysis used 27 DIPG samples and 6 brainstem LGG comparator samples, with probe-to-gene collapse and low-variance filtering performed before limma modeling. Analysis code and processed intermediate tables are available from the authors upon reasonable request.

## Ethics Statement

This study used publicly available, de-identified datasets and did not involve new human-subject recruitment, intervention, or access to identifiable private information. No additional institutional review board approval was required.

## Author Contributions

A.S. conceived the research direction, designed the computational workflow, supervised dataset selection and analysis, guided interpretation of results, and revised the manuscript. A.R. contributed to background literature review on DIPG biology, manuscript drafting, review of manuscript content, and interpretation of the mitotic hub-gene findings. S.V. contributed to dataset organization, figure generation and review, review of expression-analysis outputs, and interpretation of immune and dependency results. All authors reviewed and approved the final manuscript.

